# Degree and site of chromosomal instability define its oncogenic potential

**DOI:** 10.1101/638460

**Authors:** W.H.M. Hoevenaar, A. Janssen, A.I. Quirindongo, H. Ma, S. Klaasen, A. Teixeira, G.J.A. Offerhaus, R.H. Medema, G.J.P.L. Kops, N. Jelluma

## Abstract

Most human cancers are aneuploid, due to a chromosomal instability (CIN) phenotype. Despite being hallmarks of cancer, however, the roles of CIN and aneuploidy in tumor formation have not unequivocally emerged from animal studies and are thus still unclear. CIN can both promote and suppress tumorigenesis, but variances in mechanisms and timings of CIN induction in different oncogenic backgrounds and associated tissues limit interpretation of the contributions of CIN. Using a novel conditional mouse model for diverse degrees of CIN, we find that a particular range is sufficient to drive very early onset spontaneous adenoma formation in the intestine, showing that CIN can act as a much more potent oncogenic driver than was previously reported. In mice predisposed to intestinal cancer (*Apc*^*Min*/+^), moderate but not low CIN causes a remarkable increase in adenoma burden in the entire intestinal tract, especially in the distal colon, more closely modelling human disease. Strikingly, high levels of CIN promote adenoma formation in the distal colon even more than moderate CIN does, but have no effect in the small intestine. Our results thus show that CIN can be potently oncogenic, but that certain levels of CIN can have contrasting effects in distinct tissues.

## INTRODUCTION

Aneuploidy - an abnormal number of chromosomes - is a hallmark of human tumors^1^. While during embryogenesis almost all aneuploidies are lethal^2^ and aneuploidy levels in normal tissues are very low^3^, chromosomal aberrations are observed in 70-90% of solid tumors^4,5^. Aneuploidy is the result of chromosomal instability (CIN): the occasional gain or loss of whole chromosomes during mitosis. CIN can lead to genome destabilization^6–9^, and is associated with high intra-tumoral genomic heterogeneity, immune evasion, and promotion of metastases (reviewed in^10^). Furthermore, CIN and aneuploidy are correlated with tumor aggressiveness, therapy resistance and poor patient prognosis^11–19^.

One example of a very common and deadly cancer type that has a high prevalence of CIN and aneuploidy is colorectal cancer (CRC). CRC is divided in heritable and sporadic types, both mainly consisting of microsatellite unstable (MIN, 15%) and CIN (85%) tumors^20^. CIN correlates with poor patient prognosis: a meta-analysis of 63 studies with a total of 10.126 CRC patients (60% aneuploid tumors, as a proxy for CIN) showed that CIN tumors responded worse than non-CIN tumors to 5-Fluoruracil treatment, and (progression free) survival was decreased in CIN patients^14^.

Despite the correlations described above, the roles of CIN and aneuploidy in tumor formation are still unclear. Mouse models of CIN have occasionally shown sporadic, spontaneous tumors with very long latency (>12-18 months), and predominantly in spleen and lung^21–29^, suggesting CIN is not a potent cancer driver. In mice predisposed to cancer, CIN is either neutral^30–32^, promotes tumor formation^23,26,27,30,33–39^, or, in some conditions, suppresses it^29,30,32,37^. Comparisons between these studies is however exceedingly difficult due to the use of different oncogenic backgrounds, to differences in tissues that were examined^40^, and to the manner and time by which the tissues were exposed to CIN. Moreover, technical limitations often precluded direct measurements of CIN in the relevant tissues, and oncogenic effects in the different models could not be attributed to distinct CIN levels.

We therefore established a genetic mouse model that allows controlled induction of various degrees of CIN in a tissue-specific manner. With this model, we made direct comparisons and found striking, and tissue-type-depending, differences in the consequences for tumorigenesis between the various degrees of CIN.

## RESULTS

### An allelic series for graded increases of CIN *in vivo*

To enable tissue-specific induction of a range of CIN levels, we created mouse strains carrying a conditional T649A or D637A mutation in the spindle assembly checkpoint kinase Mps1. These mutations in human cell lines caused mild or severe CIN, respectively^41^ (Fig. 1A, B, S1A-C). We reasoned that combining these Cre-inducible *Mps1* knock-in (*CiMKi*) alleles (together and with wild-type *Mps1*) would result in an allelic series of CIN, ranging from very low (few missegregations with mostly mild errors) to very high (many missegregations with mostly severe errors).

**Figure 1.**
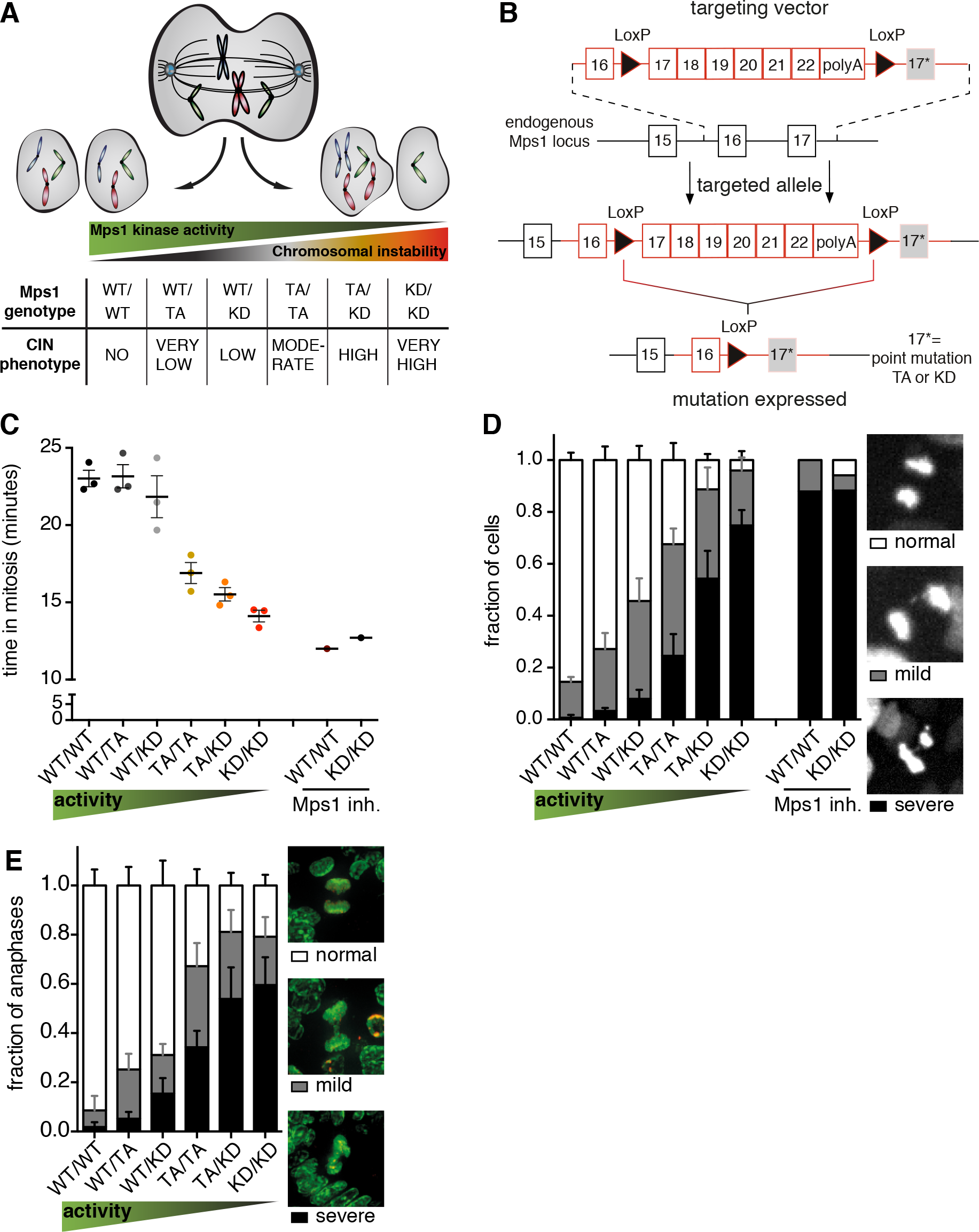
An allelic series for graded increases in CIN *in vivo*. **(A)** Scheme showing the theoretical inverse correlation of decreased function of the spindle assembly checkpoint (by genetically reduced Mps1 activities) with increased severity of CIN. **(B)** Schematic overview of generation and activation of the CiMKi alleles: the targeting vectors harbor a cDNA cassette of WT exons 17-22, that is flanked by lox-P sites, and the mutated exon 17 (TA or KD, indicated with an asterisk) in the right homologous recombination arm. In the targeted *CiMKi* alleles, wild-type *Mps1* is replaced with mutant *Mps1* upon Cre-mediated loxP recombination. **(C)** Quantification of time in mitosis (prophase to anaphase) by time lapse imaging of immortalized MEFs of the *CiMKi;Rosa26-CreERT2* genotypes 56 hours after 4-OHT addition. Mps1 inhibitor CPD5 was used as control. Error bars indicate ± SEM of three independent MEF lines per genotype; 50 cell divisions per line. See also Supplemental movies 1. **(D)** Quantification of chromosome segregation fidelity by time lapse imaging of immortalized MEFs of the *CiMKi;Rosa26-CreERT2* genotypes 56 hours after 4-OHT addition. Mps1 inhibitor CPD5 was used as control. DNA was visualized by H2B-mNeon. Segregations were scored as indicated; images depict examples. Error bars as in C. See also Supplemental movies 1. **(E)** Quantification of chromosome segregation fidelity in situ in small intestine of *CiMKi;Rosa26-CreERT2* mice one week after tamoxifen injection. DAPI (green) and anti-H3S10ph (red) were used to identify mitotic nuclei. Graph shows quantification by category as in D and E, for at least 47 anaphases per small intestine. Error bars indicate ± SD (n=3-12 mice per genotype, WT/WT group includes oil controls of other genotypes).

*CiMKi* mice were born healthy and at Mendelian ratios. Activation of Cre recombinase by addition of 4-hydroxytamoxifen (4-OHT) to mouse embryonic fibroblasts (MEFs) from *CiMKi;Rosa26-CreER*^*T2*^ mice resulted in efficient recombination and expression of mutant *Mps1* mRNA (Fig. S1D), from which mutant Mps1 protein was translated to comparable levels as wild-type protein (Fig. S1E). As expected, the allelic series caused graded reductions in Mps1 activity, as evidenced by acceleration of mitosis after mutant induction^42^ (Fig. 1C) and reduced Mad1 levels at kinetochores^43^ (Fig. S1F). Time-lapse microscopy of 4-OHT-treated immortalized MEFs showed a striking increase in mitotic errors with diminishing Mps1 kinase activities (Fig. 1D, Movies S1), verifying the predicted phenotypes of the allelic series. As expected, induced CIN resulted in increased aneuploidy in primary MEFs (Fig. S1G). Mutant induction also occurred efficiently *in vivo* in four-week old *CiMKi;Rosa26-CreRT2* mice (Fig. S1H), and analysis of anaphase figures in intestinal tissue sections showed that the expected range of CIN was induced (Fig. 1E). We thus conclude that the CiMKi mouse model enables spatio-temporal control of a range of CIN *in vivo*.

### Moderate CIN causes early onset spontaneous tumorigenesis in the intestine

Whole-body mutant inductions in *CiMKi;Rosa26-CreER*^*T2*^ mice disrupted small intestinal tissue organization (Fig. 2A). The extent of disorganization correlated with the degree of CIN and likely explained the severe weight loss seen in mice with high and very high CIN (Fig. S2A). To study early CIN induction in the intestine without adverse effects on other organs, we generated *CiMKi;Villin-Cre* mice to enable mutant induction specifically in the intestinal tract from 12.5 days post coitum (dpc)^44^. Although all *CiMKi;Villin-Cre* mice were overtly healthy (Fig. S2B), moderate CIN had caused one or more adenomas in the small intestine of two-thirds (4/6) of the mice by as early as 12 weeks of age (Fig. 2B, C). These adenomas were positive for nuclear β-catenin, showing that CIN was sufficient to induce constitutive Wnt pathway activation (Fig. 2D). To our knowledge, this is the first report of early onset, spontaneous tumorigenesis as a result of CIN. By the age of eight months, a wider range of CIN had caused increases in spontaneous adenomas (Fig. 2E, F), further underscoring a role for CIN in tumor initiation. In line with what we observed at 12 weeks, the strongest effect was again found in mice with moderate CIN.

**Figure 2:**
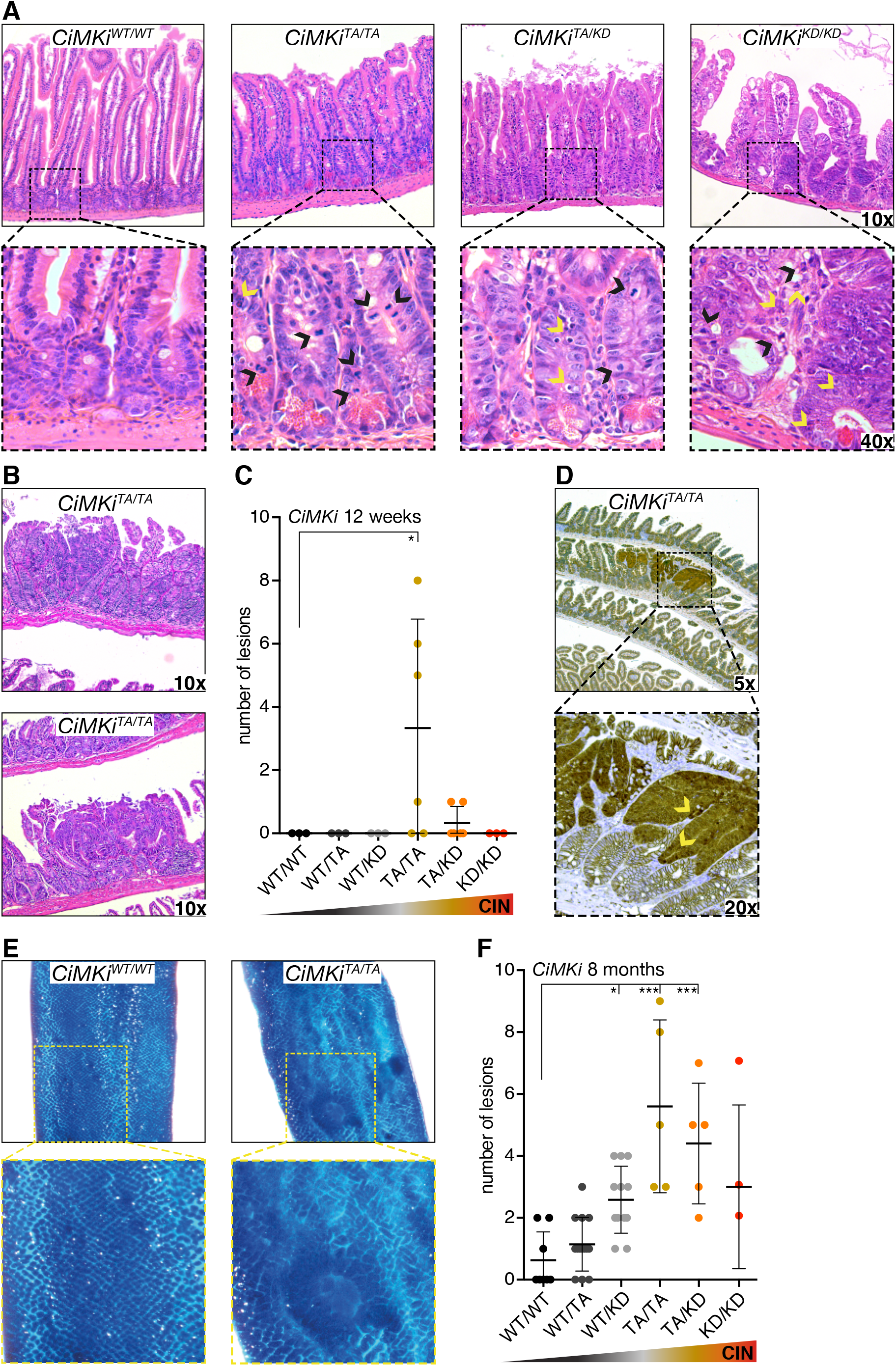
Moderate CIN causes early onset spontaneous tumorigenesis in the intestine. **(A)** H&E stained sections of *CiMKi;Rosa26CreERT2* small intestines one week after tamoxifen injection, showing aberrant crypt and cell size, hyperproliferation (black arrowheads indicate mitotic cells) and apoptotic bodies in crypt (yellow arrowheads). (B) H&E stained examples of small intestine adenomas from 12-week old *CiMKi^TA/TA^;VillinCre* mice (moderate CIN). (C) Quantification of adenomas as determined by H&E staining in small intestine tissue in of 12-week-old *CiMKi;VillinCre* mice. Data represents mean ± SD, n=3-6 per genotype. p<0.05 (*). **(D)** ß-catenin immunohistochemistry on small intestine lesions in 12-week old *CiMKi^TA/TA^;VillinCre* mice. Arrowheads indicate nuclear ß-catenin. **(E)** Examples of methylene blue stained, formalin-fixed whole mount small intestine of 8-month-old *CiMKi;VillinCre* mice. Zoom boxes indicate normal (WT/WT) and aberrant (TA/TA) mucosa. **(F)** Quantification of lesions as determined on whole mount methylene blue staining in small intestine of 8-month-old *CiMKi;VillinCre* mice. Data represents mean ± SD (N=3-14 per genotype), asterisk indicate significance (one-tailed t-test, comparing each group to WT/WT, p<0.001 (***), p<0.005 (**), p<0.05 (*)).

### Degree and site define oncogenic potential of CIN in tumor-prone intestines

Human colorectal cancers (CRCs) are often aneuploid^45^, and the vast majority is caused by loss-of-function mutations in genes of Wnt pathway components such as *APC*^46–48^. Moreover, loss of heterozygosity (LOH) of *APC* causes extensive polyp growths in patients with familial adenomatous polyposis coli (FAP) syndrome^49–51^. To examine the impact of CIN on a tissue predisposed to cancer, we next investigated mice carrying a mutant *Apc* allele (*Apc*^*Min*/+^). These mice normally develop ±30 adenomas in the small intestine and no or very few in the colon^52^. Note that the expected degrees of CIN in this tissue in *CiMKi* mice were directly verified by *in situ* analyses (see Fig. 1E). In contrast to *CiMKi;Villin-Cre* mice, very high CIN in the *Apc*^*Min*/+^ background (*CiMKi;Apc*^*Min*/+^*;Villin-Cre*) was embryonic lethal, precluding further analysis of this level of CIN. While mice with low CIN were sacrificed by the expected 12 weeks of age^30,36,53^, mice with moderate or high CIN had to be sacrificed at 6-8 weeks due to severe weight loss. *Apc*^*Min*/+^ mice with moderate CIN presented with a striking increase in the amount of small intestinal adenomas (Fig. 3A-C, S3A). Neither high nor low CIN, however, increased tumorigenesis, suggesting that adenoma formation in the small intestine is sensitive to a narrow range of CIN.

**Figure 3:**
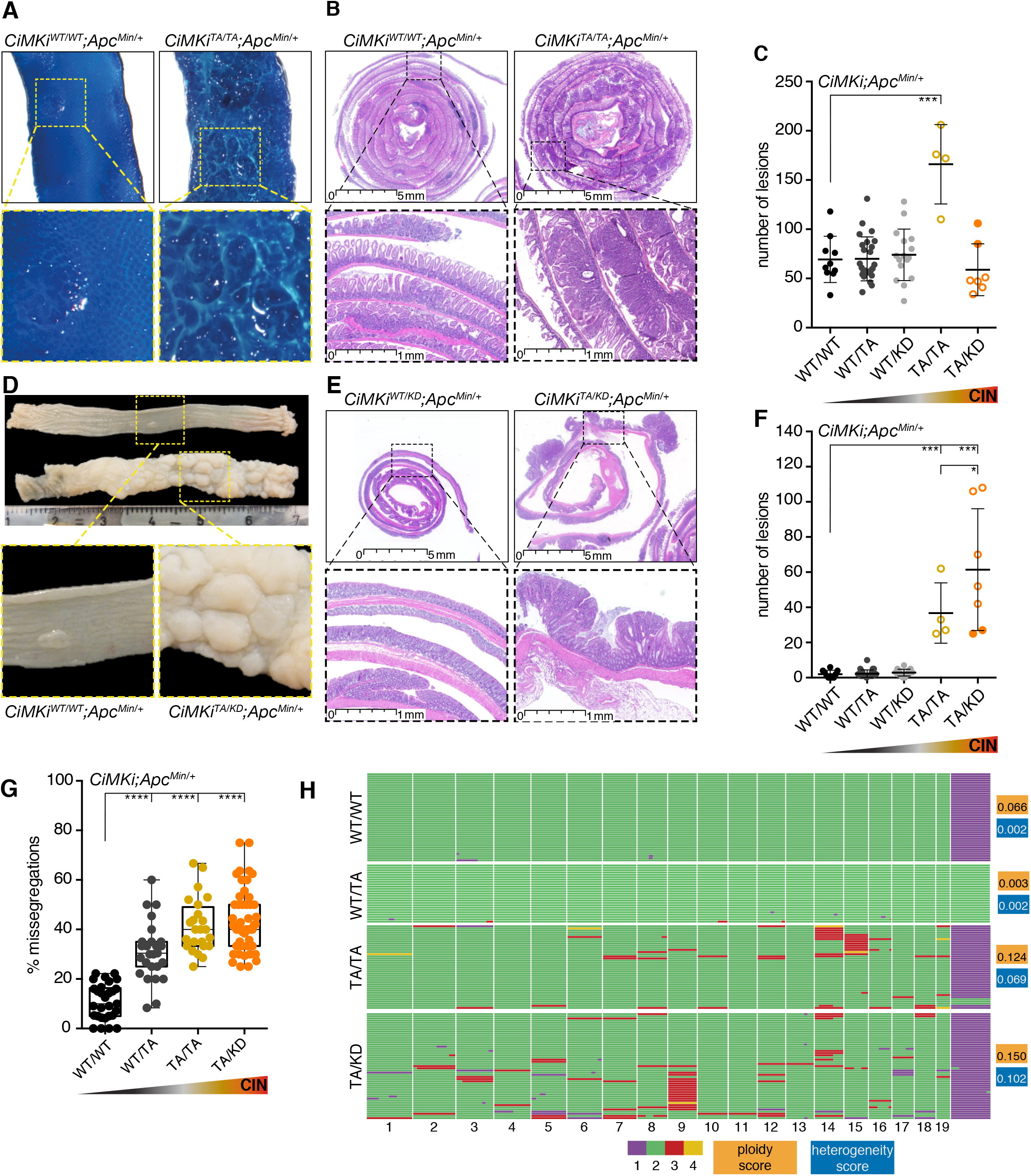
Degree and site define oncogenic potential of CIN in tumor-prone intestines. **(A)** Examples of formalin-fixed, methylene blue stained whole mount intestines of 12-week-old WT/WT and 7-week-old TA/TA mice (*Apc*^*Min*/+^*;VillinCre* background), showing mucosal architecture and abnormalities. **(B)** H&E sections of 12-week-old WT/WT and 7-week-old TA/TA mice (*Apc*^*Min*/+^*;VillinCre* background). **(C)** Quantification of small intestine adenomas on methylene blue stained whole mount small intestine of *CiMKi;Apc*^*Min*/+^*;VillinCre* mice. Each mouse is represented by an individual dot (n=4-25 mice per group), data represents mean ± SD, asterisk indicate significance (one-tailed t-test, comparing each group to WT/WT, p<0.001 (***)). Open dots represent mice euthanatized at 6-8 weeks of age, closed dots represent mice euthanatized at 12 weeks of age. **(D)** Formalin-fixed whole mount colons of 12-week-old WT/WT and 7-week-old TA/KD mice (*Apc*^*Min*/+^*;VillinCre* background) are predominantly located in the distal colon. Zooms indicate adenoma(s) in both genotypes. **(E)** H&E examples of colons of 12-week-old WT/WT and 7-week-old TA/KD mice (*Apc*^*Min*/+^*;VillinCre* background). **(F)** Quantification of adenomas on methylene blue stained whole mount colon tissue. Each mouse is represented by an individual dot (n=4-25 mice per group), data represents mean ± SD, asterisks indicate significance (one-tailed t-test, comparing each group to WT/WT, p<0.001 (***), p<0.05 (*). Open dots represent mice euthanatized at 6-8 weeks of age, closed dots represent mice euthanatized at 12 weeks of age. **(G)** Quantification of chromosome segregation fidelity by time lapse imaging of colon adenoma organoids (n=3-5 adenomas from different mice per genotype). Missegregations were quantified for at least 20 organoids with at least 5 divisions per genotype. Box-plot: each dot is one organoid, center line is median, box extends from 25th to 75th percentile, whiskers show min-to-max (one-tailed t-test, comparing each group to WT/WT, p<0.0001 (****). **(H)** Single cell whole genome karyosequencing (bin size 5 MB) showing ploidy in individual cells of three colon adenomas per genotype: aneuploidy and heterogeneity scores are given for each sample. Graph shows cells of one example per genotype. Green is 2n, purple 1n, red 3n and yellow is 4n for a given chromosome.

Macroscopic examination of the colons of the same *CiMKi;Apc*^*Min*/+^*;Villin-Cre* mice revealed that in contrast to control and low CIN mice, colons from moderate and high CIN mice were widely covered with large adenomas (Fig. 3D-F, S3D). This was most striking in the distal region of the colon, where aneuploid *APC*-mutant tumors also most frequently occur in humans^54,55^. Incidence was 100% (Fig. S3G), and the number of adenomas was substantially higher than reported for other CIN models in the *Apc*^Min/+^ background^30,36,37^. Importantly, CIN in organoids established from these colon adenomas still corresponded to the expected levels (Fig. 3G, Movies S2), suggesting that high CIN levels were not selected against after adenoma initiation and that there was no drift towards an ‘optimal’ CIN level. Of note: while individual adenoma sizes were comparable between all induced CIN levels (Fig. S3B, C, E, F), adenomas with moderate or high CIN level had reached this size substantially earlier (6-8 vs. 12 weeks). We thus hypothesize that CIN advanced initiation, accelerated growth, or both.

In humans, tumors in the distal part of the colon are often considered CIN as they are typically aneuploid and karyotypically heterogeneous^56,57^. Since the *CiMKi;Apc*^*Min*/+^*;Villin-Cre* mice with moderate and high CIN mimicked such distal colon tumors, we next assessed aneuploidy and heterogeneity of copy number alterations (CNAs) of colon adenomas. Single cell whole genome karyotype sequencing (scKaryo-seq) showed that both aneuploidy and karyotype heterogeneity were increased with moderate and high CIN (Fig. 3G, H). Chromosome 18, which harbors the *Apc* allele (that is often subject to LOH in human FAP tumors^46,52^), was diploid in the vast majority of cells. Since adenoma initiation in *Apc*^*Min*/+^ mice requires LOH of wild-type *Apc*^58^ and since CiMKi colon adenoma organoids grew independently of Wnt ligands, this indicated that LOH of *Apc* by CIN occurred in a manner other than whole chromosome 18 loss, as previously suggested^30^. Targeted PCR detected only *Apc*^Min^ alleles (Fig. S3H), strongly suggesting that LOH was accomplished either by double non-disjunction events of both chromosomes 18 or by somatic recombination^59,60^, the latter of which is likely the cause of *Apc* LOH in FAP patients^50,61^. Both these processes could be accelerated by CIN.

### Colonic crypts retain proliferating CIN cells more readily than small intestinal crypts

Our data thus far show that the effect of CIN on karyotype heterogeneity and tumorigenesis in identical genetic backgrounds depends on the degree of CIN and the tissue in which CIN occurs. As high CIN caused massive colonic adenomas but did not increase adenoma formation in the small intestine, the effects of a similar range of CIN can be profoundly different in different tissues. To better understand the tissue-dependent sensitivities to CIN, we first assessed the possibility that the same *CiMKi* mutations had resulted in different CIN levels in small intestine versus colon. Although time-lapse imaging of colon and small intestinal organoids showed that CIN levels were not identical between the two tissues (in *Apc*^*Min*/−^ background), genotypes with drastically different impacts on tumorigenesis (Fig. 3C, F) had comparable CIN levels (e.g. TA/TA in colon vs TA/KD in small intestine) (Fig. 4A, B). Strikingly however, moderate and high CIN caused significant expansion of the proliferative compartments in the colons of *CiMKi;Apc*^*Min*/+^*;Villin-Cre* mice at four weeks of age (roughly the time of adenoma initiation), but not in the small intestine (Fig. 4C, D). The percentage of proliferating cells within the compartment (proliferative index) was similar across genotypes (Fig. 4C, D, S4A-D), hence cells were more readily retained in a proliferate state in the colons of moderate and high CIN mice, increasing the chance that transformed cells propagate in colonic crypts. The fact that this increased proliferative state was not observed in the small intestine again underscores the difference in CIN response between these tissues.

**Figure 4:**
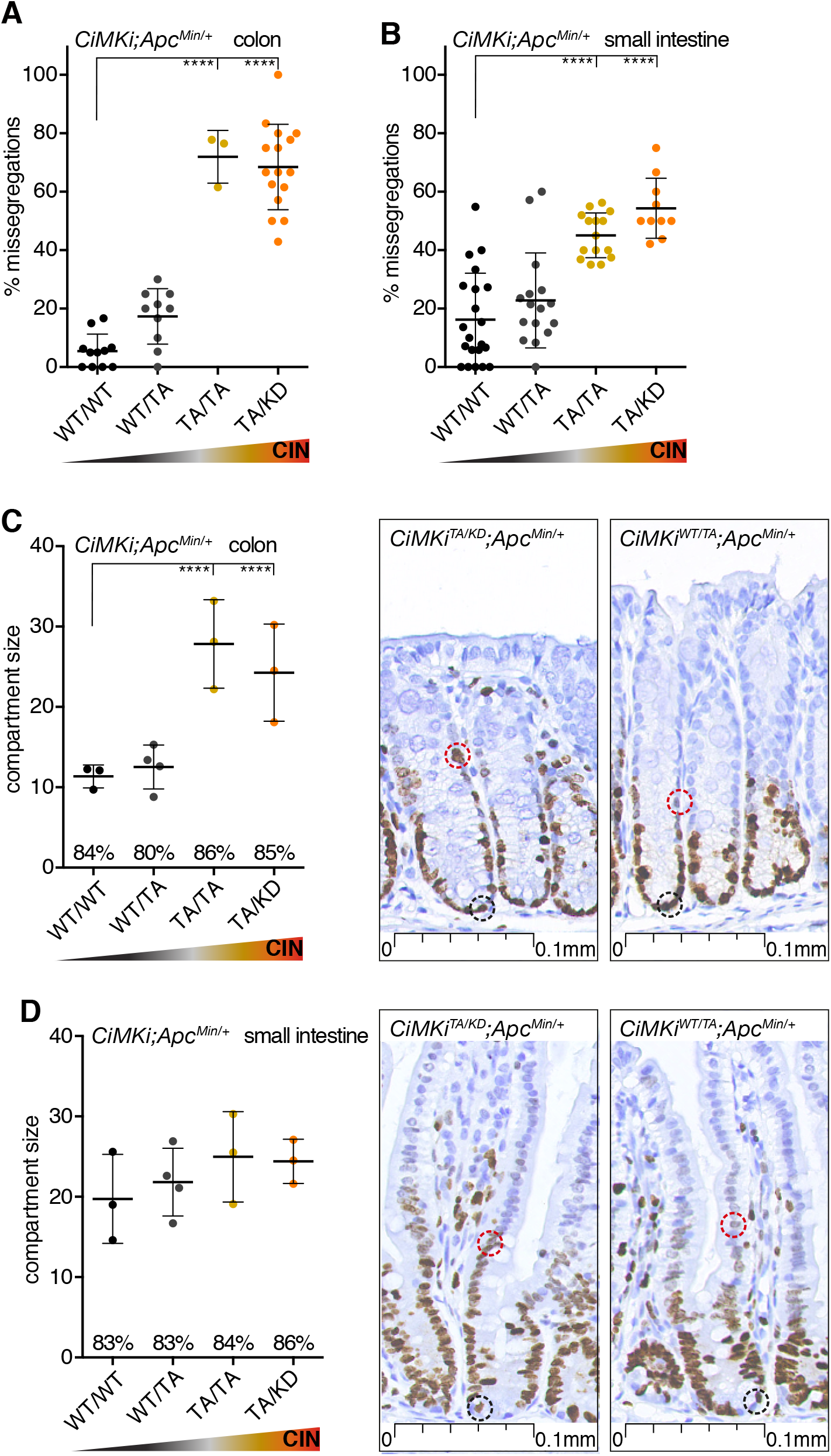
Colonic crypts retain proliferating CIN cells more readily than small intestinal crypts. **(A, B)** Quantification of chromosome segregation fidelity by time lapse imaging of colon organoids (A) and small intestine organoids (B) from various *CiMKi;Apc*^*Min*/+^*;VillinCreERT2* genotypes. Missegregations were quantified for at least 10 organoids with at least 5 divisions per line. Data represents mean ± SD, each dot is one organoid (one-tailed t-test, comparing each group to WT/WT, p<0.0001 (****). **(C, D)** Proliferative compartment in colon (C) and small intestine (D) of 4-week old *CiMKi;Apc*^*Min*/+^*;VillinCre* mice as determined on Ki-67 stained FFPE slides by scoring the number of cells between the first positive cell at the bottom of the crypt (black dotted circle) and the last positive cell in the transit amplifying zone (red dotted circle). Images show example crypts of colon and small intestine of TA/KD and WT/TA mice. Dot plot shows the average size of the compartment for each mouse (n=3-4 per genotype, 10 crypts per mouse) and error bars represent SD. Asterisk indicate significance (one-tailed t-test, p<0.01 (**), p<0.05 (*)). Percentages indicate proliferative index (percentage of Ki-67 positive cells within compartment) for each genotype.

## DISCUSSION

Aneuploidy and CIN are hallmarks of cancer, yet despite impressive efforts to model CIN in tumorigenesis, its importance to tumor development remained unclear. Our new mammalian model for inducible, graded CIN levels, combined with direct visualization of chromosome segregation error frequencies in the relevant tissues allowed us to show that 1) moderate to high CIN levels are sufficient to drive adenoma formation at early age, 2) the maximum effect on tumorigenesis is achieved by distinct CIN levels, and 3) this differs between tissues with an identical cancer predisposition mutation.

Although previous studies have reported spontaneous tumor formation in CIN mice, it occurred sporadically and with late onsets of more than 12 months of age^21–29^. By contrast, our CiMKi mice with moderate CIN (TA/TA) developed a substantial number of lesions in the small intestine as early as 12 weeks of age. This shows that CIN is a more potent driver of tumorigenesis than previously thought. As our model was able to probe the effects of a wide range of CIN, it is possible that the optimal CIN level for tumor induction was not reached in prior studies. The inducible nature of the CiMKi model is most probably a prerequisite for reaching the higher CIN levels. Many other CIN models were not inducible or tissue-specific, resulting in embryonic lethality in homozygous knock-outs, possibly due to severe missegregations during the developmental stages^22,23,62–64^. Also, we used the CiMKi model here to induce CIN locally in the gastro-intestinal tract, thereby preventing adverse effects in other tissues.

Our model enabled us to directly compare the effects of the various CIN levels between two different tissues -small intestine and colon- within the same mice. Whereas in the *Apc*^*Min*/+^ background moderate CIN levels markedly increased adenoma burdens in both tissues, a higher CIN level did not affect the small intestine, but increased the adenoma burden in the colon even more. Our data therefore do not support the previously proposed model in which low CIN levels promote tumorigenesis while high CIN leads to cell death and tumor suppression^65,66^. Instead, we argue that the role of different CIN levels is much more complicated, as certain levels of CIN can have contrasting effects in distinct tissues.

There might be several explanations for the different effects between small intestine and colon: our finding that moderate and high CIN leads to an enhanced proliferative state in colon but not in small intestine shows a remarkable difference in response between the two tissues. Enhanced proliferation can increase the chance that transformed cells propagate in colonic crypts^67^, however it does not account for the tumorigenic effect of moderate CIN in the small intestine. Therefore, other factors might play a role as well, such as the normal variations throughout the gut in stem cell number and physiological Wnt activity^68^ or in the adaptive immune landscape^69^. It will be exciting to further investigate the underlying mechanisms, as it can impact on future treatment strategies for different cancer types. The tissue-specific inducible nature of the CiMKi model enables studying the impact of various CIN levels in many other organs as well.

*Apc*^*Min*/+^ mice^53,70^ have been widely used to model human FAP, the hereditable form of CRC^51^. Tumorigenesis in these mice requires LOH of *Apc*, which directly mimics the human disorder, and is genetically comparable to tumorigenesis in FAP patients. However, whereas in human FAP patients mostly colon tumors occur, *Apc*^*Min*/+^ mice develop many adenomas in the small intestine but very few in the colon. It is therefore intriguing that addition of moderate to high CIN to this model causes early occurrence of colon adenomas, thus making it a better model for human disease. Furthermore, as in human FAP, colonic adenomas of *CiMKi;Apc*^*Min*/+^ mice are predominantly located in the distal colon and are aneuploid and karyotypically heterogeneous^71^. Also, in contrast to adenomas from *Apc*^*Min*/+^ mice^58^, LOH in *CiMKi;Apc*^*Min*/+^ mice did not occur by loss of (part of) the wild-type allele. Instead, we found disomy of the mutant chromosome 18 in the adenomas, similar to a CIN model driven by Bub1 insufficiency^30^. Other processes by which LOH can be achieved are somatic recombination events^60^, as was described for *APC* LOH in human FAP tumors^50,61^, or double non-disjunction events during mitosis^30^. CIN can be involved in both processes: DNA damage and double strand breaks as a result of CIN may be repaired through recombination with the mutant allele, and doubling of the mutant chromosome accompanied or followed by loss of the wild-type allele by non-disjunctions during anaphase can lead to disomy of the mutant. So even though the exact mechanism remains to be uncovered, LOH of *Apc* in the *CiMKi;Apc*^*Min*/+^ mice is most probably accelerated by CIN. Taken together, *CiMKi;Apc*^*Min*/+^ mice are a useful new model to study sporadic and hereditary human CRC.

In conclusion, it is now possible with the CiMKi model to accurately study the interaction between CIN and tumor development in a host of tissues and genetic backgrounds. Because of tight spatio-temporal control of CIN, CiMKi also enables investigations into the effect of various levels of CIN on cancer cell dissemination, as well as on possible tumor regression. The latter may greatly aid ongoing efforts that examine if exacerbating CIN, for example by Mps1 small molecule inhibitors, has potential as cancer therapy.

## ACKNOWLEDGMENTS

We thank H. Snippert for helping designing the CiMKi targeting vectors, J. Jonkers for advice and sharing reagents, and H. Clevers for sharing reagents and help with ES cell targeting. We thank S. van Mil and J. van Rheenen for donating mouse strains. We thank the Hubrecht mouse facility, the Hubrecht FACS facility, the Hubrecht Imaging facility, Single Cell Discoveries, and the Utrecht Sequencing Facility (USEQ) for assistance with the experiments. This work is part of the Cancer Genomics Centre and the Oncode Institute, and was further supported by the Dutch Cancer Society (grant numbers HUBR-2012-5321, HUBR-2012-5513, and 10126).

## AUTHOR CONTRIBUTIONS

W.H.M.H., A.J., R.H.M., N.J. and G.J.P.L.K. designed the research. W.H.M.H., A.J., and N.J. analyzed and assembled the data. A.I.Q. assisted with animal maintenance and experiments. H.M. and G.J.A.O. performed pathology analyses. S.K. performed and analyzed single cell karyosequencing. A.T. assisted with organoid culture and experiments. W.H.M.H., N.J. and G.J.P.L.K. wrote the paper.

## METHODS

### Mice: strains, experiments and analysis

All animal experiments were approved by Animal Experimental Committee and the Dutch Central Authority for Scientific Procedures on Animals (CCD). All animals were bred and housed under standard conditions at the animal facility of the Gemeenschappelijk Dieren Laboratorium (GDL), Utrecht, the Netherlands, and the Hubrecht animal facility. Genetically modified mice strains used in this study include: *ACTB:FLPe* (B6.Cg-Tg(<u>ACTFLPe</u>)9205Dym/J, stock number 005703), *Rosa26-CreER*^*T2*^ (B6.129Gt(ROSA)26Sor^tm1(cre/ERT2)Tyj^/J, stock number 008463), and *Apc*^*Min*/+^ (C57BL/6J-Apc^Min^/J, stock number 002020) and were purchased from JAX® mice. *Villin-Cre* mice we a gift from S. van Mil (originated from JAX® mice (B6.Cg-Tg(Vil1-cre)997Gum/J, stock number 004586). *Villin-creER*^*T2*^ mice were a gift from J. van Rheenen. All mice were maintained in C57BL/6 background. Mice were genotyped using standard PCR and targeted sequencing procedures. For primers see Table 1.

*CiMKi* mice were generated under license of UMCU (DEC 2010.I.02.026). The CiMKi alleles were designed following the example of the BRAF-V600E inducible model by Dankort et al.^72^: a cDNA cassette of wild-type exons 17-22 is followed by a stop codon and polyA sequence. The cassette is flanked by loxP sites, and the 3’ recombination arm harbors one of the point mutations in exon 17 (D637A: GAT>GCT; T649A: GCA>ACA). The cassette was cloned into the pAC16 targeting vector (kind gift from J. Jonkers). For a detailed outline of construction of CiMKi vectors see Supplementary Information.

129/Ola-derived IB10 ES cells (kind gift from H. Clevers) were electroporated (Biorad gene pulser) with the linearized targeting construct pAC16-D637A or -T649A. Targeted cells were selected with puromycin (1μg/ml, Sigma) and single colonies were subsequently picked and cultured in 96-wells plates.

Individual clones were analyzed for presence of the CiMKi alleles using standard PCR (Primers used: Forward TCTATGGCTTCTGAGGCGG and Reverse AAGGGACATCAGGGAAGCAA). DNA from targeted ES cells yielded a band of ~2.8 kb. Southern blot was performed according to standard protocols to confirm correct integration of the CiMKi alleles. 5’ probe (500bp) was obtained from genomic DNA from 129/Ola-derived IB10 ES cells, and labelled using a standard Rediprime II Random Prime labelling system (GE healthcare) and radioactive [α-32P] dCTP. Digestion of genomic DNA from ES cells with EcoRV and hybridization with the 5’ probe resulted in a 9.5 kb band (when wildtype) or 4.7 kb band (when targeted with CiMKi allele).

Confirmed targeted ES cell clones for both CiMKi mutations were injected into C57BL/6 blastocyst, which were then transplanted into pseudo-pregnant females (standard techniques, performed under the license of the GDL Utrecht). Chimeric mice were bred with C57BL/6 mice to obtain germline transmission. Agouti mice were then backcrossed six times into a C57BL/6 background. Genotypic analysis of offspring was performed using standard PCR and targeted sequencing (Table 1, supplementary information). To remove the puro cassette from the original pAC16 construct, CiMKi mice were bred with ACT-Flp mice (C57BL/6 background). Only lines that showed loss of the puro cassette (as confirmed by standard PCR) were used to maintain CiMKi lines.

To induce loxP recombination, MEFs were treated with 4-hydroxy-tamoxifen (4-OHT; 1μM, Sigma H6278). Mice were injected intraperitoneally with Tamoxifen (1 mg dissolved in corn oil; Sigma, C8267). In *CiMKi;Apc*^*Min*^*;Villin-Cre* mice, CiMKi alleles were induced at 12.5 dpc. when the Villin promotor is activated. To confirm recombination, RNA was isolated with a quick RNA kit (Zymo Research). cDNA was prepared using standard procedures, subjected to PCR and subsequently sequenced to determine the presence of T649A or D637A. For primers see Table 1, supplementary information.

Mice were sacrificed at four weeks, twelve weeks or eight months of age, and immediately dissected. small intestine was separated from colon, both were flushed with PBS and pieces of tissue were snap-frozen for later RNA/protein analysis. The organs were stored in formalin until further processing.

### Histology and Immunohistochemistry

Formalin fixed intestines were cut open longitudinally and stained with 0.25% methylene blue in dH20, and rinsed with PBS. Pictures were taken with 6.3x magnification using an Olympus SZX stereo microscope to count the number of lesions in the small intestine and colon. After washing with PBS to remove the methylene blue, intestines were rolled into “Swiss rolls” for paraffin embedding.

For identification and assessment of lesions 4-μm sections of paraffin-embedded tissue were cut and stained with hematoxylin/eosin (H&E). These slides were scanned (Nanozoomer XR, Hamamatsu) for digital image analysis. Grading of dysplasias was done following the existing guidelines for human intestinal adenomas.

Apoptotic bodies were recognized on H&E stained sections, according to strict morphological criteria such as cell shrinkage with retracted pink to orange cytoplasm, chromatin condensation and nuclear fragmentation and separation of cells by a halo from adjacent enterocytes.

For proliferation measurements slides were incubated with Ki-67 antibody (ThermoFisher, RM-9106, ARS pH9, 1:50 and 3-hour incubation). Sections were counterstained with hematoxylin, dehydrated and coverslipped using Pertex. Ki-67 positive cells were counted from the bottom to the top of the crypt till the upper most positive cell. The proliferative compartment was defined as the part of the crypt between the bottom and the upper most labelled cell. The proliferative activity (Ki-67 index) was calculated as the percentage of positively labelled cells divided by the total number of counted cells within the proliferative compartment. ß-catenin localization was assessed on paraffin sections stained with anti-ß-catenin (BD Transduction Laboratories, Clone 14/Beta-Catenin 610154, 1:1000, overnight incubation), anti-Mouse Envision-HRP (DAKO, one hour) as secondary antibody, and counterstained with hematoxylin.

### MEFs (isolation, immunofluorescence, mitotic spreads, Western blot, and live microscopy)

CiMKi mice were bred with *Rosa26-CreER*^*T2*^ mice and maintained in a stable homozygous *CreER*^*T2*^ background. Pregnant females were sacrificed at 13-17 dpc. by cervical dislocation. Uterine horns were dissected out and placed in tubes containing PBS. Embryos were separated from their placenta and surrounding membranes. Red organs, brains and tail (for genotyping) were removed. Embryos were finely minced using razor blades and the remaining cells/tissues were suspended in a tube containing 2 ml Trypsin and kept at 37°C for 15 minutes. Two volumes of media (DMEM supplemented with 10% FBS, non-essential amino acids, glutamin and Pen/Strep) were added and remaining tissues were removed by allowing them to settle down at the bottom of the tube. Supernatant was subjected to centrifugation for 5 minutes at 1000 rpm, cell pellet was resuspended in medium and plated in 10 cm dishes.

For immunofluorescence cells were plated on 12mm coverslips and harvested after one hour nocodazole (250 ng/ml, Sigma, M1404) and MG132 (2 μM, Sigma C2211) treatment. Cells were pre-extracted with 0.1% Triton X-100 in PEM (100 mM PIPES (pH 6.8), 1 mM MgCl_2_ and 5 mM EGTA) for one minute at 37°C before fixation with 4% paraformaldehyde in PBS. Coverslips were subjected to antibody staining following standard procedures (primary antibodies anti-Mad1 (Santa Cruz sc67337, 1:1000), anti-Centromere Protein (ACA) (Antibodies Incorporated 15-234-0001, 1:2000). Images were acquired on a DeltaVision RT system (Applied Precision) with a ×100/1.40NA UPlanSApo objective (Olympus) using SoftWorx software. Images are maximum intensity projections of deconvolved stacks. Quantifications were done using ImageJ software and a macro to threshold and specifically select kinetochores as described previously^73^.

For mitotic spreads, MEFs were treated with STLC (1 μM, Sigma 164739) for 4 hours. Mitotic cells (isolated by shake-off) were treated for 10 minutes in hypotonic buffer (75 mM KCl), fixed with acetic acid/methanol, dropped onto glass cover slides and stained with DAPI (1mg/ml, Sigma 32670). Images were acquired on a DeltaVision RT system (Applied Precision) with a ×100/1.40NA UPlanSApo objective (Olympus) using SoftWorx software. Chromosomes were counted manually using Image J software.

For Western blot, MEFs were treated with 4-OHT or EtOH for 72 hours, and then lysed with Laemmli buffer. Protein levels were assessed by standard Western blot procedures (anti-ESK (Santa Cruz sc-541, 1:1000), and anti-α-Tubulin (Sigma T5168; 1:10000)).

For live cell imaging, immortalized MEFs (transduced with large T and small T expressing lentivirus (Plasmid #22298, Addgene) were transduced with an H2B-Neon expressing lentivirus (pLV-H2B-Neon-ires-Puromycin)^74,75^, and selected with puromycin (1μg/ml, Sigma P7255). These stably H2B-mNeon expressing MEFs were plated in 24-well plates and imaged 56 hours after 4-OHT treatment for sixteen hours in a heated chamber (37°C and 5% CO2) using a ×20/0.5NA UPLFLN objective on an Olympus IX-81 microscope, controlled by Cell-M software (Olympus). Images were acquired using a Hamamatsu ORCA-ER camera and processed using Cell-M and ImageJ software.

### Organoids (isolation, culture and live microscopy)

Organoids were isolated from *CiMKi;Apc*^*Min*/+^*;VillinCre(ERT2*) mice as described previously^76^. In brief, intestines of six-to-twelve weeks-old mice were dissected and cleaned with PBS. They were incubated in 0.5mM EDTA on ice for 30 minutes (normal tissue), or EDTA treatment followed by 45 minutes in DMEM 2% FBS 1% Pen/Strep supplemented with 75u/ml collagenase and 125 μg/ml Dispase (adenoma tissue). Intestines were put in tubes with PSB, and crypts were removed from their niche by harsh shaking. After filtering the suspension using a 70 μm strainer, crypts or tumor cells were seeded in Matrigel (Corning, 356231). Organoids were cultured in medium containing advanced DMEM/F12 medium (Invitrogen,126334-010), Hepes Buffer (Sigma, H0887,1 mM), Pencilin/Strep (Sigma, P0781, 1%), Ala-Glu (Sigma, G8541, 0.2 mM), R-Spondin conditioned medium (20%, kind gift from Hans Clevers) (wild-type only), Noggin conditioned medium (10%) (Thermo/Life Technologies, PHC1506, 1x), B27 (Thermo/Life Technologies, 17504001, 1x), nicotinamide (Sigma-Aldrich, 72340, 10 mM) (colon only), N-acetylcysteine (Sigma-Aldrich, A7250, 1.25 mM), EGF 0,1% (Invitrogen/Life Technologies, 53003-018) and Primocin 0.5% (Invivogen, ant-pm1). For passaging, organoids were sheared by repetitive pipetting and re-plated in Matrigel in a pre-warmed 24-well plate.

To establish stable organoid lines expressing H2B-mNeon, organoids were transduced with an H2B-Neon expressing lentivirus (pLV-H2B-Neon-ires-Blasticidin)^74,75^, and selected with blasticidin (InvivoGen; 20μg/ml). For induction of CiMKi alleles organoids were treated with 1 μM 4-OHT for 56 hours. Organoids were seeded and imaged in 8-chamber IBIDI slides using a confocal spinning disk (Nikon/Andor CSU-W1 with Borealis illumination), equipped with atmospheric and temperature control. Organoids were imaged in XYZT-mode (12 to 20 z-sections at 2.5 μm intervals, for 8 to 12 hours) at 37°C at 3-minute intervals, using a 30X silicon objective and an additional 1.5X lens in front of the CCD-camera. 3% 448nm laser and 50nm disk pinhole were used. Raw data were converted to videos using an ImageJ macro as described^74,77^. Fidelity of all observed chromosome segregations was scored manually, guided by custom-made ImageJ/Fiji macro for ordered data output.

### Single cell karyotype sequencing (scKaryo-seq)

Snap-frozen adenoma tissue was stained with 10 μg/ml Hoechst 34580 (Sigma-Aldrich) and minced in a petri dish, on ice, using a cross-hatching motion with two scalpels. The minced tissue was kept on ice for 1 hour after which it was filtered through 70 μm and 35 μm strainer. Nuclei were sorted in a 384-well plate containing 5 μl of mineral oil (Sigma) in each well and stored at −20°C until further processing for library preparation and sequencing, as previously described^78,79^. Libraries were sequenced on an Illumina Nextseq 500 with 1 x 75 bp single-end sequencing. The fastq files were mapped to GRCH38 using the Burrows-Wheeler Aligner. The mapped data was further analyzed using custom scripts in Python, which parsed for library barcodes, removed reads without a NlaIII sequence and removed PCR-duplicated reads. Copy number analysis was performed as described previously^80^. The data are deposited in the ENA repository (accession number PRJEB31573).

### Statistical analysis

Power analysis predicted the number of animals that had to be used in each group to detect differences with 80% power and 95% confidence. Animals were not randomized, but assigned to the experimental groups based on their genotype. Statistical analyses were done with GraphPad Prism software. Comparisons between CiMKi wildtype and CiMKi mutants were analyzed with one-tailed student’s t-tests. Data is presented as mean ± SD unless otherwise stated in legends.

