## Supplementary Information for "Degree and site of chromosomal instability define its oncogenic potential"

**Table 1: Primers for genotyping and cDNA analysis**

| **Gene** | **Forward primer** | **Reverse primer** | **Expected band size** | **Sequence primer** | **Mutation sequence** |
|---|---|---|---|---|---|
| *CiMKi* | *CCAAATGGCTAGGGGAGCCACTGATG* | *GGTGAGGTTGTTTCCAACTGGTAG* | **Mutant 250 bp** | NA | NA |
| *CiMKi* (mutation) | GTGTCCTCACCCTGAAAATG | CAAAGCACAGCTGGGCTGTAGAG | **~1000 bp** | CGGATTTTATTTTGAAGGTATTG | T649A ACAA/GCA D637A TTGGA/CT |
| *CiMKi* cDNA (mutation) | CCTAGAAGACGCCGATAGCC | GTCTCTGATTGCTTCTGGGGC | **~400 bp** | GATAAGATCATCCGCCTCTATG | T649A ACAA/GCA D637A TTGGA/CT |
| *Apc$^{Min/+}$* | *CCGGAGTAAGCAGAGACACAAG* | *CTGTCGTCTGCCACACAATG* | **~400 bp** | *CATGACTGTTCTTTCACC* | *GACAGAAGTTT/A* |
| *Rosa26-CreER$^{T2}$* (mutant) | GGCAGGAAGCACTTGCTCTCCC | CCTGATCCTGGCAATTTCG | **~825 bp** | NA | NA |
| *Rosa26-CreER$^{T2}$* (wild-type) | GGCAGGAAGCACTTGCTCTCCC | GGAGCGGGAGAAATGGATATG | **~650 bp** | NA | NA |
| Villin-Cre(ER$^{T2}$) | *CAAGCCTGGCTCGACGGCC* | *CCTGATCCTGGCAATTTCG* | **~220 bp** | NA | NA |

### Supplementary Movie Legends

**Movies S1: Increased missegregation rates in *CiMKi;Rosa26-CreER$^{T2}$* MEFs.**

**(A)** Time lapse imaging of *CiMKi$^{WT/WT}$;R26CreER$^{T2}$* immortalized MEFs expressing H2B-mNeon, 56 hours after 4-OHT addition. **(B)** As **A**, but for *CiMKi$^{KD/KD}$;R26CreER$^{T2}$*.

**Movies S2: Increase in missegregation rates in *CiMKi;Apc$^{Min/+}$;Villin-Cre* colon adenoma organoids.**

**(A)** Time lapse imaging of *CiMKi$^{WT/WT}$;Apc$^{Min/+}$;Villin-Cre* colon adenoma organoids. Color dept-coding (purple is bottom of organoid, red is top) was used to identify the position of the cells, left panel) and maximum projections are depicted in the right panel. **(B)** As **A**, but for *CiMKi$^{TA/TA}$;Apc$^{Min/+}$;Villin-Cre* colon adenoma organoids.

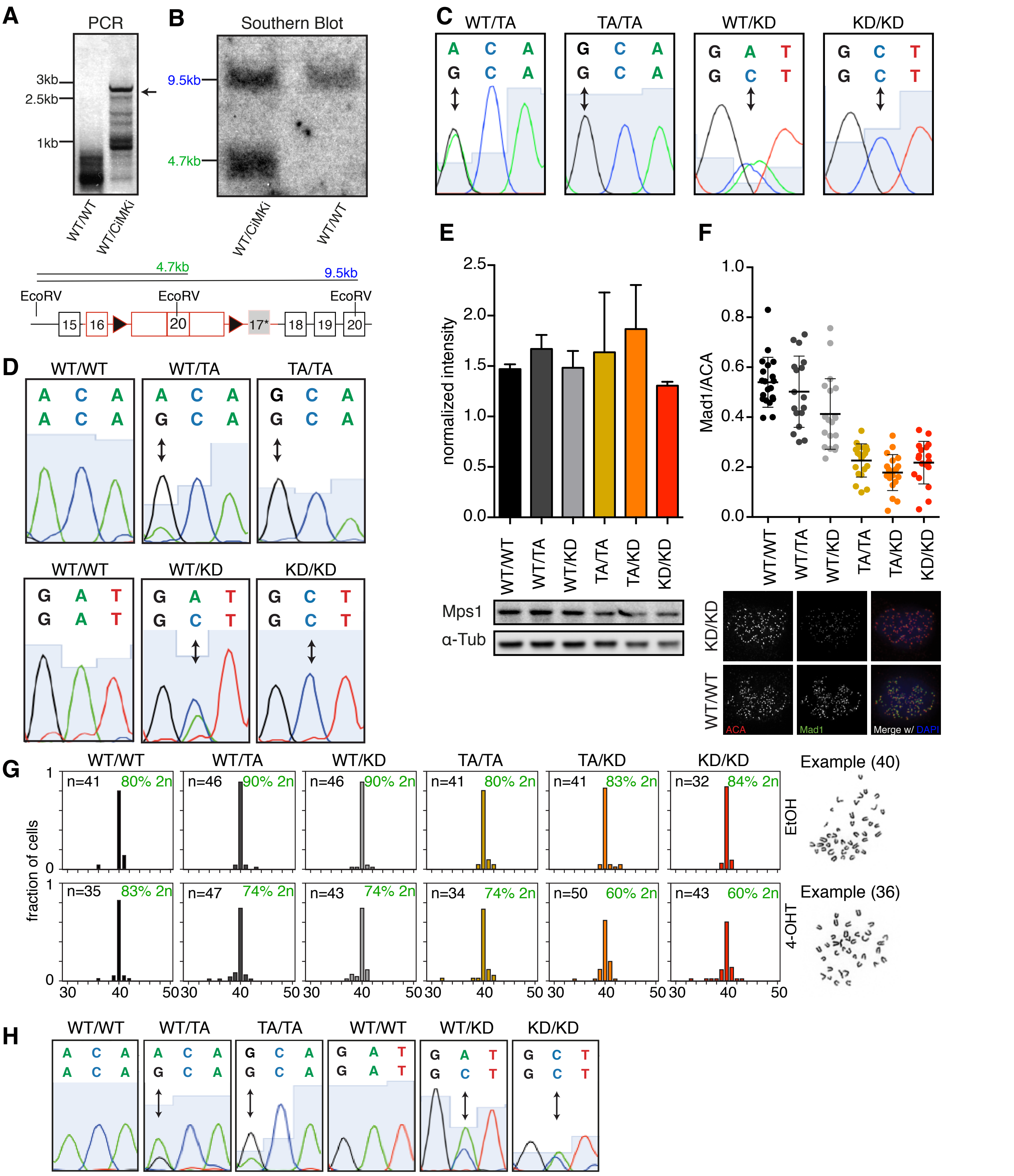


**Figure S1: A novel mouse model for CIN: Cre-inducible Mps1 Knock-in (CiMKi).**

**(A)** Genomic PCR of targeted ES cells confirming presence of the CiMKi allele. Shown here specifically is the CiMKi-T649A ES clone that was used for blastocyst injection. **(B)** Confirmation of correct integration of the CiMKi alleles by Southern blot. Shown here specifically is the CiMKi-T649A ES clone that was used for blastocyst injection. Lower schematic shows EcoRV restriction sites used for Southern blot, indicated 5' of exon 15 and in exon 20. **(C)** Targeted sequencing confirming presence of Mps1 mutations in mouse ear genomic DNA. **(D)** RT-PCR followed by targeted sequencing of cDNA from CiMKi MEF lines 56 hours after 4-OHT addition shows hetero- or homozygous expression of both D637A (A to C) or T649A (A to G) alleles. **(E)** Western blot of Mps1 protein expression in *CiMKi;Rosa26-CreERT2* MEFs 56 hours after 4-OHT addition. Intensity is normalized over α-tubulin. **(F)** Examples and quantification of Mad1 localization on kinetochores as a proxy for Mps1 activity in *CiMKi;Rosa26-CreERT2* MEFs 72 hours after 4-OHT addition. Cells were blocked in mitosis by nocodazole and MG132 for 30 minutes. Graph shows quantifications of kinetochore signals as ratios over ACA signals. Data represents mean ± SD of at least 20 cells per condition. **(G)** Examples and quantification of diploid and aneuploid cells on metaphase spreads (DAPI) of *CiMKi;R26CreERT2* primary MEFS 56 hours after 4-OHT addition. MEFS were blocked in mitosis by 4 hours treatment with nocodazole. Ploidy was assessed by counting the number of chromosomes per cell, percentage of diploid cells is given. **(H)** RT-PCR followed by targeted sequencing on cDNA from *CiMKi;R26CreERT2* small intestine tissue one week after tamoxifen injection confirms effective recombination and expression of the mutant alleles. Hetero- or homozygous expressions of both D637A (A to C) or T649A (A to G) alleles are shown.

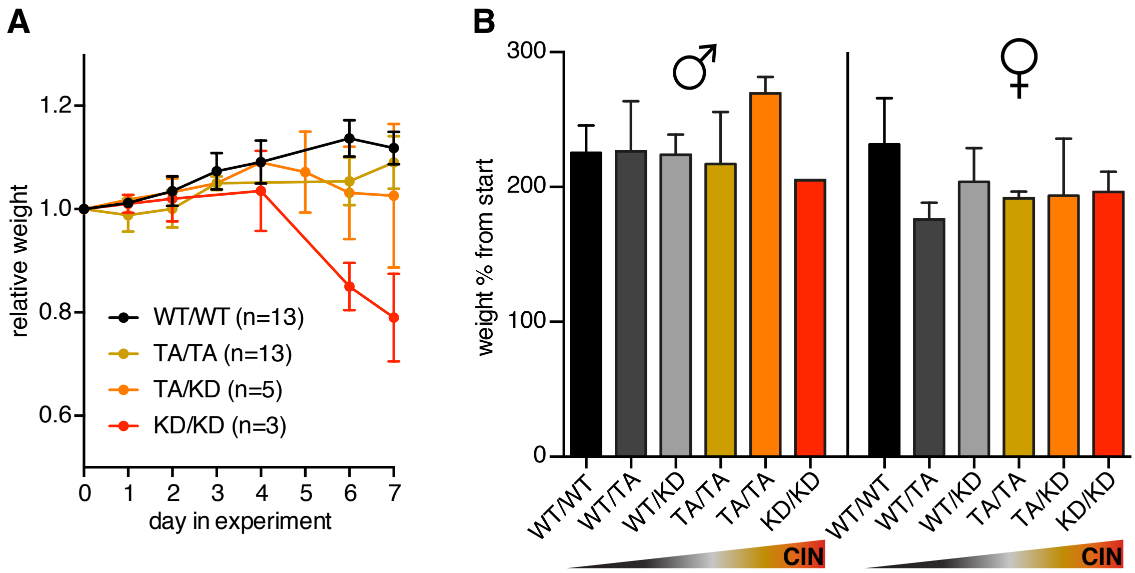


**Figure S2: CIN leads to spontaneous tumorigenesis in the intestine.**
**(A)** Relative bodyweight of *CiMKi;R26CreERT2* mice after three consecutive days of intraperitoneal tamoxifen injection. Lines represent change in bodyweight per group as fraction of their weight at the start of the experiment (mean ± SD). **(B)** Relative body weight in male (left) and female (right) *CiMKi;VillinCre* mice of all genotypes. Increase in weight is shown as percentage from start of the experiment (4 weeks) to end (8 months). Data is shown as mean percentage increase ± SD.

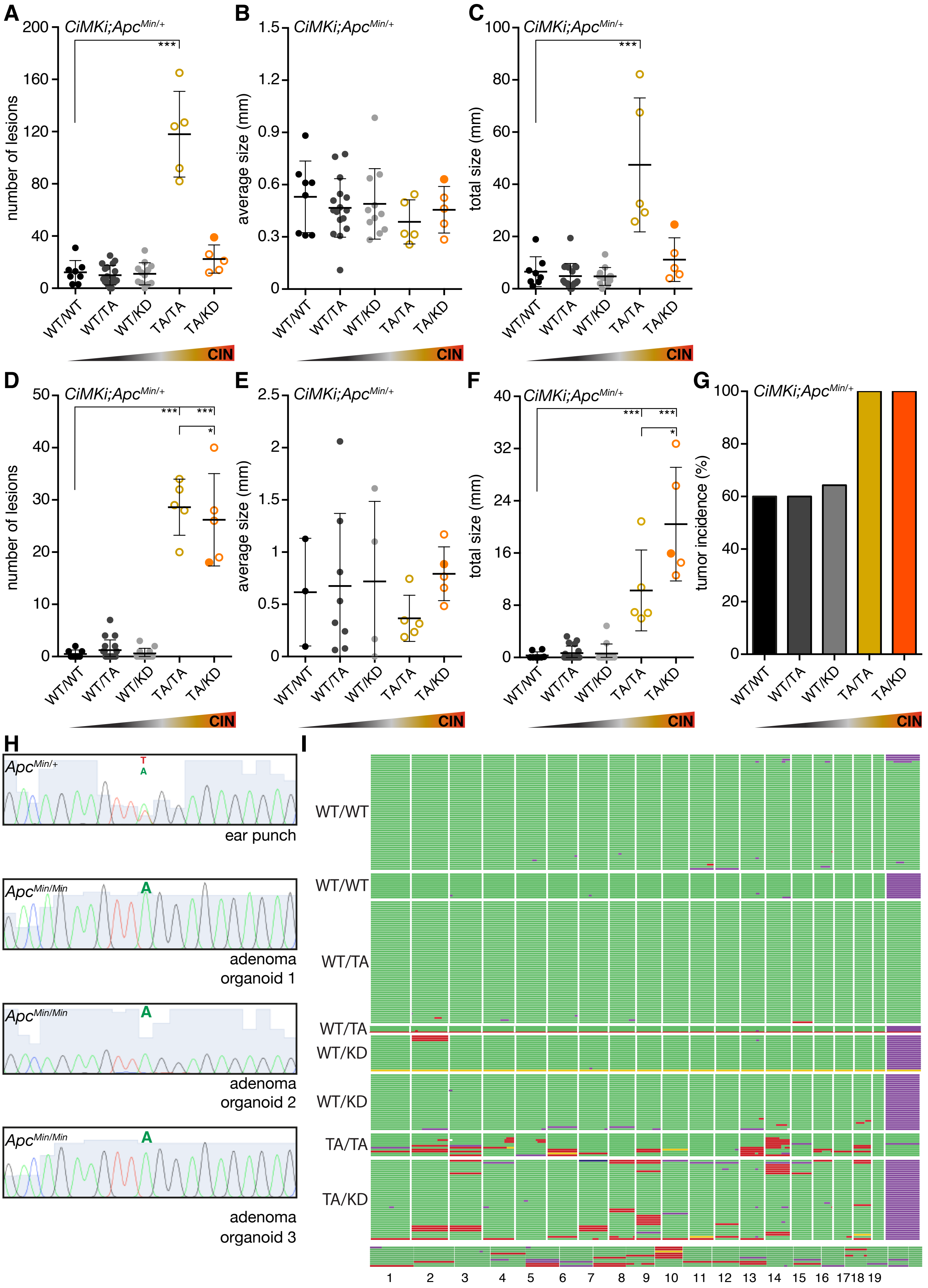
A
B
C
D
E
F
G
H
I
CiMKi;ApcMin/+
number of lesions
average size (mm)
total size (mm)
tumor incidence (%)
WT/WT
WT/TA
WT/KD
TA/TA
TA/KD
CIN
***
*
ApcMin/+
ear punch
ApcMin/Min
adenoma organoid 1
adenoma organoid 2
adenoma organoid 3
T
A
1 2 3 4 5 6 7 8 9 10 11 12 13 14 15 16 17 18 19

**Figure S3. CIN differently affects small intestine and colon adenoma formation in *$Apc^{Min/+}$* mice.**

**(A)** Quantification of small intestine adenomas on H&E sections of *CiMKi;$Apc^{Min/+;}$VillinCre* mice. Each mouse is represented by an individual dot (n=4-15 mice per group), data represents mean ± SD, asterisk indicate significance (one-tailed t-test, comparing each group to WT/WT, p<0.001 (***)). Open dots represent mice euthanatized at 6-8 weeks of age, closed dots represent mice euthanatized at 12 weeks of age. **(B)** Average size of small intestine adenoma for each mouse was measured by taking the diameter of the lesion on H&E slides of *CiMKi;$Apc^{Min/+}$;VillinCre* mice. Data represents mean ± SD. **(C)** Total adenoma burden in small intestine as the sum of all adenoma diameters per mouse. Data represents mean ± SD, asterisk indicate significance (one-tailed t-test, p<0.001 (***).

**(D)** Quantification of colon adenomas on H&E sections of *CiMKi;$Apc^{Min/+}$;VillinCre* mice. Each mouse is represented by an individual dot (n=4-15 mice per group), data represents mean ± SD, asterisk indicate significance (one-tailed t-test, comparing each group to wild-type, p<0.001 (***), p<0.05 (*)). Open dots represent mice euthanatized at 6-8 weeks of age, closed dots represent mice euthanatized at 12 weeks of age. **(E)** Average size of colon adenoma for each mouse was measured by taking the diameter of the lesion on H&E slides of *CiMKi;$Apc^{Min/+}$;VillinCre* mice. Data represents mean ± SD. **(F)** Total adenoma burden in colon as the sum of all adenoma diameters per mouse. Data represents mean ± SD, asterisk indicate significance (one-tailed t-test, p<0.001 (***), p<0.05 (*)). **(G)** Colon adenoma incidence in *CiMKi;$Apc^{Min/+}$;VillinCre* mice of the indicated genotypes. **(H)** Targeted sequencing confirming absence of wild-type *Apc* genomic DNA in adenoma organoids. *$Apc^{Min/+}$* mouse ear genomic DNA is given as reference. **(I)** Single cell whole genome karyosequencing (bin size 5 MB), showing cells of two examples per genotype (see also Fig. 3H). Green is 2n, purple 1n, red 3n and yellow is 4n for a given chromosome.

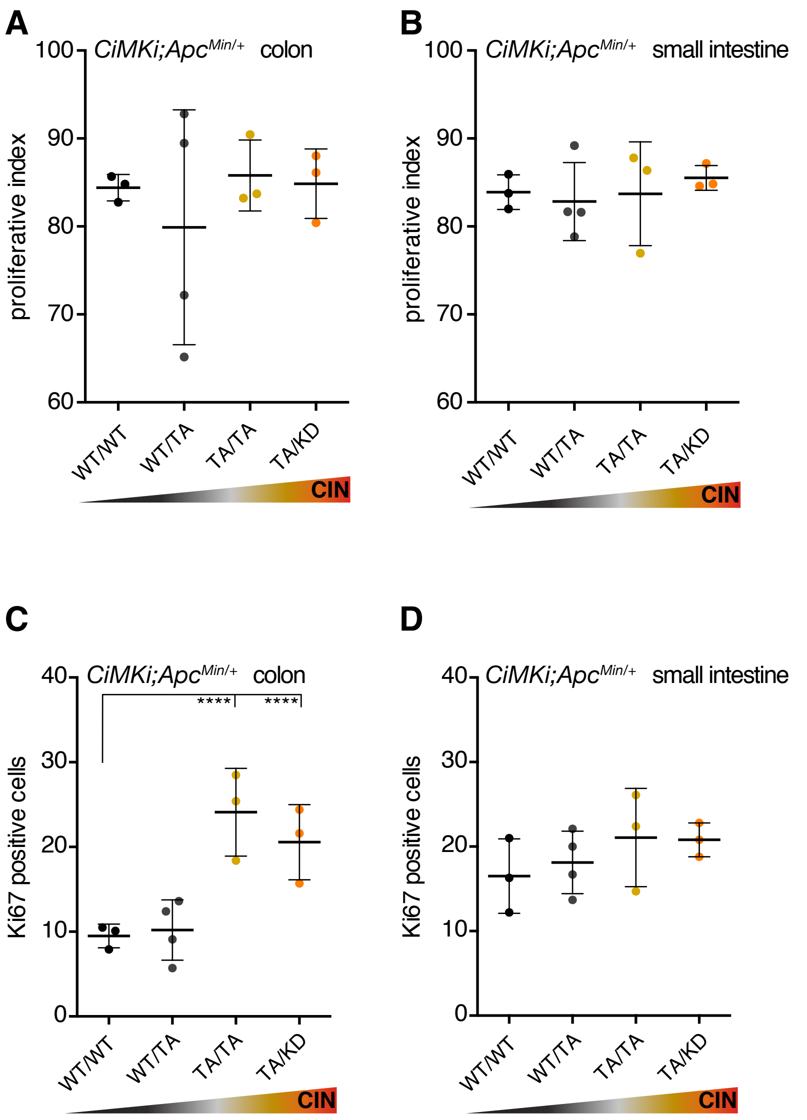


**Figure S4: Colonic crypts retain proliferating CIN cells more readily than small intestinal crypts**
**(A, B)** Proliferative index in colon (A) and small intestine (B) of 4-week old CiMKi;Apc$^{Min/+}$;VillinCre mice as determined on Ki-67 stained FFPE slides by calculating the percentage of Ki-67 positive cells within the proliferative compartment. Data represents mean ± SD, n=3-4 mice per genotype. **(C, D)** Number of Ki-67 positive cells in proliferative compartment I colon (C) and small intestine (D) of 4-week old *CiMKi;Apc$^{Min/+}$;VillinCre* mice. Data represents mean ± SD, n=3-4 mice per genotype. Asterisk indicate significance (one-tailed t-test, p<0.01 (**), p<0.05 (*)).

### Extended Methods

*Cloning of CiMKi targeting vectors*

CiMKi alleles were cloned into the pAC targeting vector (based on pFlexible[1]) (kind gift from Jos Jonkers). Three fragments were ligated separately into the vector as follows:

1) Conditional fragment into HindIII-PacI (between loxP sites)
2) 5’ Recombination arm into PmeI-AscI (upstream of 5’ loxP site)
3) 3’ Recombination arm, including either point mutation, into SbfI-NotI (downstream of 3’ loxP site).

Details for the three fragments:

*1) Conditional fragment*

The conditional fragment was obtained by assembling three separate PCR products by ligation into pCDNA3 (see details below):

#1=*HindIII*-(last part of)intron16-*XhoI* (~600 bp)

#2=*XhoI*-**exon17-exon18-exon19-exon20-exon21-exon22**-*BamHI* (~1000 bp)

#3=*BamHI*-intron22(polyAsignal)-*PacI* (~600 bp)

This resulted in the following complete sequence of the conditional fragment (exon17-22 part (PCR product #2) in bold):

*AAGCTT*ATGGCCAGTTTTTCAGTCTCCAAAGGATTTTCTCTTTGGAGCCCGGTGTGAGAGT
TGGTCTGTTCTTGCTGTTTTCTGTTTCATTATTTGTTTGTTCTTGTTCTCTCCAGTTTCCTCT
GCCTTTCCCTTGTCTTTTAAACAGATGTGTTCAATCTCATGCGCTTCCTCTGCTTCTCCTTT
TAGAAGCAGTAACTAAGCAGAGGATAGGACTCTTTGGGGGTGAGGGAGGCCTTGCGATC
TTATTGTCTGACAGTTCTTGGTGGTGCTTTCCTGGCCTGTGCATGGCAGTGTGCAGCACAC
ACTGACAGGCTAGGCCTGGAATCCTGAACAGTTCACTTCAACTCAAATCACATTGAGTGT
CCTCACCCTGAAAATGATGATGATAATGGGACTTATTTGCTATGCATTGCTATAGAATGG
TACCTGGCCCACAACATTTGCTAAGAAAGTGTAGAATGTATAATAACTAAATATTAATGT
GTTGCCTAATTGAAAGGATAAGCCTGACCTAAAGATTATGAAATGCTTTTTTGTCCCCTG
GCCATAGAATGTTTTTCTTCATCTAACAGAACGGATTTTATTTTGAAG**GTATTGTTCATA**
**GTGATCTGAAGCCTGCTAACTTTGTGATAGTGGATGGAATGCTAAAGCTAATTGATT**
**TTGGGATTGCAAACCAAATGCAGCCAGACACAACAAGCATTGTTAAAGATTCTCAG**
**GTTGGCACAGTTAACTATATGGCCCCAGAAGCAATCAGAGACATGTCTTCTTCAAG**
**AGAAAATTCGAAAATCAGAACCAAGGTAAGTCCCAGAAGTGATGTCTGGTCCTTGG**
**GGTGCATTTTGTACTACATGACTTATGGGAGGACGCCATTTCAGCACATCATCAATC**
**AGGTCTCTAAACTGCACGCCATAATCAACCCTGCTCATGAGATTGAATTTCCCGAGA**
**TTTCGGAAAAAGATCTTCGAGACGTGCTAAAGTGCTGTTTAGTGAGGAACCCTAAA**
**GAGAGGATATCTATCCCTGAGCTCCTCACACATCCGTATGTTCAAATTCAGCCCCAT**
**CCAGGCAGCCAAATGGCTAGGGGAGCCACTGATGAAATGAAATATGTGTTGGGTCA**

**ACTTGTTGGTCTGAATTCTCCTAACTCCATCTTGAAAACTGCAAAAACTTTGTATGA
ACGTTATAATTGTGGTGAAGGTCAAGATTCTTCGTCATCCAAGACTTTTGACAAAAA
GAGAGAAAGAAAGTGATGCACAGCTACGTACAAACCAAGAACACTAGATTGTTTCC
TCTGCCATACTCTTGAATCTCTGAGGAAATCTACCAGTTGGAAACAACCTCACCTGG
ATTTTATCAGTTAAAAAAACAAACAAACAAAACTTCAGTAGATTATCCTCAAAAGAA
AGCTGTAAAGTTAACCACTCATAGCACTGTGTATATTAAATTATAGAGTTGTGCTTT
TCTTTTATGCTTTTCTGTAAATCTGCTAATGTTTTACGTTTGGAACAGTGAATGATA
GCTGGAATGTTGAAGAGCTCTGTAAATAAAGCGTCACCACAGTTCC**AGAACTGTACA
GTGGTCAGTTTCTTCAATCAAATGTGTTCTTGGCATGATAGCAAAATTTTTAGAAAAACG
GGATTAAGAATAGACCGTAGTAAATAAAGTTTAACAATTAAATTTCCAAAGGATTTAGG
ACTGTAAACAGGCTCACACCTTTAGTCCCAGCACTTGGGAGGCAGAGGCAGGCAGATTT
CTGAGTTCGAGGCCAGCCTGGTCTACAGAGTGAGTTCCAGGACAGCCAGGGCTACACAG
AGAAACCCTGTCTTGGCGGGGAGGGGCCGGGGAGGGGGGGAGATCCAAAGGATGTAGG
AGGCAGAGACAGGTGGATCTCTGTGATTCTGAAGCCAGCCTGGTCTACAGAACTAGTTCT
AGGATAGCCAAGGCTATACAGACACTTTCTCCCCACCACCATCCTGCCCTCAAAAAAATT
GTAAATAAATTTCCTAATTGTGTACAGCCATGATACCATCTATAGTATTTGGTCTGCAAG
TGGCTTTTTCAGTTCTCCCTTTGACTCTTCAAAGTACATATGGGGTTTGGTGTTCTAAATA
TTGTGCTGGAGTTTGTGATTTAATGTCTATAGTTAATACATGCCATTATTGAG*TTAATTAA*

Product #1 was obtained by standard PCR using genomic DNA from 129/Ola-derived IB10 ES cells (kind gift from Hans Clevers). Primers used (*Italic* is overhang, in *Capitals/Italic* the restriction sites):

#1 Forward: *cggcgAAGCTT*ATGGCCAGTTTTTCAG

#1 Reverse: *cgccgCTCGAG*CTTCAAAATAAAATCCGTC

Product #2 was obtained by standard PCR using mouse cDNA (Imagenes, Germany, BC 058851, ID: 30023533). Primers used (*Italic* is overhang, in *Capitals/Italic* the restriction sites):

#2 Forward: *ccggcCTCGAGgcgctactctggtg*GTATTGTTCATAGTG

#2 Reverse: *ccgcgGGATCCgcgctactctggtg*CTGGAACTGTGGTGAC

Product #3 was obtained by standard PCR using genomic DNA from 129/Ola-derived IB10 ES cells (kind gift from Hans Clevers). Primers used (*Italic* is overhang, in *Capitals/Italic* the restriction sites:

#3 Forward: *ccgccGGATCC*AACTGTACAGTGGTC

#3 Reverse: *ccgccgTTAATTAA*CTCAATAATGGCATG

First, products #1 and #2 were ligated by standard methods into pCDNA3 simultaneously. XhoI site was removed using site-directed mutagenesis. Primers used:

(XhoI)loop Forward: GACGGATTTTATTTTGAAGGTATTGTTCATAGTGATC

(XhoI)loop Reverse: GATCACTATGAACAATACCTTCAAAATAAAATCCGTC

Second, product #3 was ligated into the pCDNA containing #1 and #2. BamHI site was removed using site-directed mutagenesis. Primers used:

(BamHI)loop Forward: GCGTCACCACAGTTCCAGAACTGTACAGTGGTCAG

(BamHI)loop Reverse: CTGACCACTGTACAGTTCTGGAACTGTGGTGACGC

This resulted is completion of the conditional fragment. Third, the conditional fragment was digested from pCDNA and ligated into pAC16 (using *HindIII and PacI restriction sites*).

*2) 5' Recombination arm*

The 5' recombination arm fragment was obtained by standard PCR using genomic DNA (from 129/Ola-derived IB10 ES cells (kind gift from Hans Clevers). Primers used (*Italic* is overhang, in *Capitals/Italic* the restriction sites):

5'arm Forward: c*tagcgGTTTAAAC*TCGAAGGCCTCAACCTCACAGAGATCTTTC

5'arm Reverse: *atcttaGGCGCGCCGGGCCC*TCTCCTCCTATCTGTAGGATG

This resulted in the following complete sequence of the 5' recombination arm:

(*PmeI*-(lastpartof)intron14-**exon15**-intron15-**exon16**-(firstpartof)intron16-*ApaI*-*AscI* (~2kb):

*GTTTAAAC*TCGAAGGCCTCAACCTCACAGAGATCTTTCTGCTTCTGCCTCTCTCCTGAGTG
CTGAGATTAAAGGTGTGTGCAGCCATGCCTAGCTGTGTGTGGCTTTCCTGTTACCACATT
CTGTGGATAGCTTATCTGTTGTTTGTGCCCCACCTTGTACTTAATATCAAGATGAGATGTT
TGGCTGTTCGGTGATACTAGCTCTGGTGACTAAAACTGAATAGGACCTTTATTCCTTTGCT
GAGTGTTCCTCCTTGGCTATCAGGCTGTTTTATGTTATGGTGATACATGAACAGAGATAA
AGGGCTTATTTTAAGTTTTACTGTAATTCTCTAAGCCAGCTAGCTGTAATTAGACTTGCTT
CTGGCTATTTTATTAACTTATAGCAAGTAATTGGAAAGCATCCCATCAGACCCATTTGTA
TAATTCCTGCACACAGTACGCTGAGGCGGGATGATTGCCTGGGCTACATACATGCACTGT
ATAGGCTGCTGCTATGGCTTTGGTGCAGGGCCCTGGCTTTAATCTCTTAACATAAAAACC
ATAAGCAAAAGACAAAATAGGTAGGAGTGTATATTTCCACATGGAGCATGTCTTCCCAT
AAATATTTTCCTTTTCACGCTCCCCTTATTAGATTTTCAGTTATGAGCATAAGGAAGAAGG
TGGGGTAAGAGTGATTGAACAAGAGTGACAGGGAAGAGGATTCGTGCAGGGGAGGGAA
GTGCATGCATGTTCTAGGCACTGAGTGATGGGTGTGCTGGAAGCTGTAAACTGCGTGGG
GGCCTCTTCCTCAGTGCTTTAAGAAATTGATTCATAGGAACATCATTGCTCCTGCCAACCT
AACTCAACTGTGACTTGCGCTGCTTTCCACAAATGAATGTAGTGATGGCTTACAATTACT
GTGATTTTTAAAAATATTCCCTATCAGAGAAATGAATTGGTTGATAGTAGGCACAATGAA
AAGGTGGGAGTTGGTGGGGAGAGGGGTCTAGGAAGTGAACTGTCAAATCACAGCCTTAG
TCATCCTTATCGTCTTCGAATGTCTTTTCCTTGATGTTTTCTCCCTGGATAAGAAAGGCAT
CCCTAGAATTTTGTGGATATAGCAACATCATATTTAAGTTGGTTTTCTTAGACACTGATGT
AGAAAACCTTTGAATTATTTGAATGTCCATTGTTATAGGGGCTGGAAATGGATACTTAGC
TTCTCATGTTGGTATTTCTTTAGGAATATCCTCAGCCTGAGACTGTTAGTGTTAAATGGAA
AGGTACTGCTCCAGTTTTCAGAGGGAGACATGTCCTAAGCTCTTTCTCCACTTTTTATGTA
G**GTGTTTCAGGTATTGAATGAGAAAAAACAGATAAACGCTATCAAATATGTGAACCT**
**AGAAGACGCCGATAGCCAAACTATTGAGAGCTACCGCAACGAGATAGCGTTTTTGA**
**ACAAACTACAGCAACACAGTGATAAGATCATCCGCCTCTATGATTA**GTATGAATTCA
TTTTTATTTTAAAAATAAAAGTTTGTTCTTGCCATAATTCTTAGGCAAAGAGTAAATCCTT
AATGACATAATGTGGGCATTTATTGTTTTGTTGTGTCTGTTTATCTTTAATTGCAG**TGAAA**
**TCACCGAGCAGTACATCTACATGGTAATGGAATGTGGAAACATTGACCTAAATAGT**
**TGGCTTAAAAAGAAAAAATCCATCAATCCATGGGAACGCAAGAGCTACTGGAAAAA**
**CATGTTGGAGGCAGTACACATAATCCATCAGCATG**GTATTTTCATATCTCTTCATACA
CGTAAAGTTAAAATAGTTGTTAATTGTGCCATTTTAGAAACATACCCTTAACTGGAAGTT

CATTAGAGGTGAAGGCACTCTTAAGAGTGGTTATACACAGGCTACAGAACACAAACAAG
CACAGGATGTAGAACAGAAATGGCCACATGTACAATGTAAACTTACCCTCCTCTGGTACC
TGGGGATTCCTATCTTCAAGTCCTGAGGATTTGGACATCCTACAGATAGGAGGAGA*GGG*
*CCCGGCGCGCC*

This fragment was ligated into pAC16 containing the conditional fragment (using *PmeI and AscI restriction sites*, upstream of 5' loxP site).

*3) 3' Recombination arm*

The 3' recombination arm fragment was obtained by standard PCR using genomic DNA (from 129/Ola-derived IB10 ES cells (kind gift from Hans Clevers)). Primers used (*Italic* is overhang, in *Capitals/Italic* the restriction sites):

3'arm Forward: *ccgccCCTGCAGG*ATGGCCAGTTTTTCAG

3'arm Reverse: *ctagcgGCGGCCGC*CTATTTGCAAATCACAAAG

This fragment was ligated into pAC16 (using *SbfI and NotI restriction sites*, downstream of 3' loxP site). CiMKi point mutations were introduced using site-directed mutagenesis.

Primers used for T649A mutation:

mMps1-T649A-F: CAAATGCAGCCAGACACA***GCA***AGCATTGTTAAAGATTC

mMps1-T649A-R: GAATCTTTAACAATGCTTGCTGTGTCTGGCTGCATTTG

Primers used for D637A mutation:

mMps1-D637A(KD)-F: GAATGCTAAAGCTAATT***GCT***TTTGGGATTGCAAAC

mMps1-D637A(KD)-R: GTTTGCAATCCCAAAAGCAATTAGCTTTAGCATTC

This resulted in the following complete sequence of the 3' recombination arm (*SbfI*-(lastpartof)intron16-**exon17***-intron17-*NotI* (~2.2kb), containing either T649A or D637A point mutation in exon 17):

*CCTGCAGG*ATGGCCAGTTTTTCAGTCTCCAAAGGATTTTCTCTTTGGAGCCCGGTGTGAGA
GTTGGTCTGTTCTTGCTGTTTTCTGTTTCATTATTTGTTTGTTCTTGTTCTCTCCAGTTTCCT
CTGCCTTTCCCTTGTCTTTTAAACAGATGTGTTCAATCTCATGCGCTTCCTCTGCTTCTCCT
TTTAGAAGCAGTAACTAAGCAGAGGATAGGACTCTTTGGGGGTGAGGGAGGCCTTGCGA
TCTTATTGTCTGACAGTTCTTGGTGGTGCTTTCCTGGCCTGTGCATGGCAGTGTGCAGCAC
ACACTGACAGGCTAGGCCTGGAATCCTGAACAGTTCACTTCAACTCAAATCACATTGAGT
GTCCTCACCCTGAAAATGATGATGATAATGGGACTTATTTGCTATGCATTGCTATAGAAT
GGTACCTGGCCCACAACATTTGCTAAGAAAGTGTAGAATGTATAATAACTAAATATTAAT
GTGTTGCCTAATTGAAAGGATAAGCCTGACCTAAAGATTATGAAATGCTTTTTTGTCCCC
TGGCCATAGAATGTTTTTCTTCATCTAACAGAACGGATTTTATTTTGAAG**GTATTGTTCA**
**TAGTGATCTGAAGCCTGCTAACTTTGTGATAGTGGATGGAATGCTAAAGCTAATT*G**
**CT*TTTGGGATTGCAAACCAAATGCAGCCAGACACA*GCA*AGCATTGTTAAAGATT**
**CTCAG**GTAGGAGTTTTGCTGTCTTGGTTGTATTTTAGTGTTTTGAACCAGGGTTTTGCATC
AGGGTTTTGCATAACCTAGAATGCTCTTGACTTTGATCAGTGGCCTTCAGCTCCTGATCCT

GCTGCCTGTGCATCCCAGGTGTGGGCTTATAGGTGTCAGCCCCGACACCCGACTTCAGGT
AGGATTTTAATGATGGCTGGTTACTACAAGGCTTAGTTCATTTTTATCTGTTAAATATGTT
GCCAATATTATATTTTTACCAACCATGTTATTCCAAAAATTTGAAGTCTTTTTAAAGAATA
GAAACTATGTTTATAAAAGACCATGGTCAAAGCCATGGTCAATTTGATTTATAAAAGCAG
TTCAAGATCAGACAAGTATATTTATGAATTTTGGATGATTTTCTCATAGCTGAGGCAGGG
CAGAGAGTAATTGCACCTTCATGTTCTCCACTGTCCTGTTTCTTTTTCTTACTGCTTAAATT
TGGGAGAAAGTTTTAAGAGAGCCTTATTGGGAATACTGAAGCGTTCCACTCAGCTACGC
GTTAAAAAGGAAATATTTTACTTACTGTTTGGGGGGCGGTGTGCGGATTCCAGATGTAAG
TGTGCCATTGTGGAGGTGTGGGGATGGCTTCGGTAAACCACTTCTCTCCTACTGTGGGCC
CTGGGAGCCAAAGTCAGATTGTGTGTGCAGCAAGCTCTACAGCCCAGCTGTGCTTTGTAG
TAACATTTGCTGTGGTAAATCTCATGAAGCTGAAGTAGTGAGGGGAAAACAGAGCTGAA
AGGTGATGTCGACTGCACCTCGCAGGCTGTGTCCAGGGATGGAGATAAATCAGAAGATA
AATTACCATGCACGTAGAAAGTCATTCTTCTTGACAGCCATTCATTGTTTTTTGTTTCAGA
AGTACAGATGATGAACAGTGAGTGTAGATGAGACTGAAGTTTTCTATGGCAAGGTCTTA
GCAGGCCGACATTTTGTTACCTTAGAACTAAAGGATTTTGCGTATTATCTCCATGCCCAG
CTAGAGAAGCTCTTCTCTGATATAGGTTTCCAACCCATCTTGATCTGCACATGGAGCCGA
AGAATATTGGGAGATAAAGCTAGCTGGTTCCTTTTATTCATGTATTAATTTGTTGCTTGGT
TTATTGAGGGAAGAATATGTTGAATTTATTGGAACATGAAAGTGAATGAAAGGCCAAGT
TCAGAATCCGCCTACGCAGTTGTAAAGACTTAGTACTTAGTACTTAGTACTTAGCACTAG
CTCTCCAGCACAGCTGCAGACAGCACAGTGCTCCCTGTGCTCCAGACGGAGCCCGTTCAT
TCTCAGCCCAGCTCATCTGATTGTACCTGGGATGGGATAGTACATACATTCTTATATTGTT
AGCAGTTATTTGAATTTTTCAAGTCTGTCATTTAAATCATTAGTTATTCAAATTTCCAAGA
ATCTGACATTTACATATTTACAAATCTAGAAAGATATTCTCATTGATTTCTTTGTGATTTG
CAAATAG*GCGGCCGC*
**(*GCT* =D637A, *GCA* =T649A)**

These fragments were ligated separately into pAC16 containing the conditional fragment and the 5’ recombination arm (using *SbfI and NotI restriction sites*, downstream of 3’ loxP site), to obtain the two separate complete targeting vectors containing CiMKi-T649A or CiMKi-D637A.

Fidelity of all PCR products, site-directed mutagenesis steps, and ligation steps were verified by sequencing.
